# Neural decoding of competitive decision-making in Rock-Paper-Scissors

**DOI:** 10.1101/2025.01.09.632285

**Authors:** Denise Moerel, Tijl Grootswagers, Jessica L. L. Chin, Francesca Ciardo, Patti Nijhuis, Genevieve L. Quek, Sophie Smit, Manuel Varlet

## Abstract

Social interactions are fundamental to daily life, yet social neuroscience research has often studied individuals’ brains in isolation. Hyperscanning, the simultaneous recording of neural data from multiple participants, enables real-time investigation of social processes by examining multiple brains while they interact. Previous hyperscanning research has largely focused on cooperative tasks, with fewer studies examining competitive contexts. Here, we obtained electroencephalography (EEG) hyperscanning data from 62 participants (31 pairs) who played a computerised version of the Rock-Paper-Scissors game, a classic paradigm for studying competitive decision-making. Although the optimal strategy is to be unpredictable and thus act randomly, participants exhibited behavioural biases, deviating from this ideal. Using multivariate decoding methods to measure neural representations within the two players’ brains in interaction, we found information about decisions made by participants during gameplay, revealing certain strategies. Notably, losers uniquely represented information about prior trials, suggesting this may impair optimal performance. These results reveal how competitive decision-making is shaped by cognitive biases and previous outcomes, highlighting the difficulty of achieving randomness in strategic contexts. This work advances our understanding of decision-making and cognitive dynamics in competitive interactions.

## 1. INTRODUCTION

Social interactions form a crucial part of our daily lives. Although studies in the field of social neuroscience have predominantly focused on individual participants during passive tasks, for example observing facial expressions and interactive behaviours (Krumhuber et al., 2023; McMahon & Isik, 2023), there has been a recent shift towards multi-brain or hyperscanning methods as they allow for the study of neural processes underlying real-time social interactions (Lehmann et al., 2023; Redcay & Schilbach, 2019). In this approach, neural data are recorded simultaneously from two or more participants interacting in real-time, enabling investigations of how brains dynamically respond and coordinate with one another (Czeszumski et al., 2020; Dumas et al., 2010; Zamm et al., 2024). Previous hyperscanning work has advanced our understanding of human cooperative activities in which brains constantly need to monitor and predict one’s own and others’ actions (Cheng et al., 2024; Dumas et al., 2010; Keller et al., 2014; Moreau et al., 2024; Tognoli et al., 2007; Varlet et al., 2020; Zamm et al., 2024). However, less research has investigated competitive activities (Babiloni et al., 2007; Balconi & Vanutelli, 2017; Cui et al., 2012; Nam et al., 2020; Sinha et al., 2016). Importantly, whereas cooperation requires behavioural predictability to facilitate the anticipation of each other’s actions and intentions, competition often requires behavioural unpredictability to gain an advantage (Glover & Dixon, 2017; Vesper et al., 2011). The current study investigates the neural processes underpinning decision performance in competitive activities.

Previous hyperscanning research has primarily focused on interbrain synchrony – the alignment of neural activity between interacting individuals – which has been shown to scale up with the level of cooperation (Fallani et al., 2010; Hu et al., 2018; Toppi et al., 2016). A few studies that directly compared interbrain synchrony in cooperative and competitive contexts have suggested that competition might be at the very end of this continuum, with evidence for stronger interbrain synchrony during cooperation compared to competition found using EEG (Chuang & Hsu, 2023; Sinha et al., 2016) and (functional) Near-Infrared Spectroscopy (fNIRS) (Cui et al., 2012; Lu et al., 2019). For example, Sinha et al., 2016 used a computerised pong game to show increased interbrain synchrony when participants played as a team against the computer program, compared to when participants played against one another (Sinha et al., 2016). The authors suggested that the difference in interbrain synchrony could in part be explained by participants having different strategies to defeat the opponent in the competition condition. Another study revealed interbrain coupling during cooperation and interbrain decoupling during competition when comparing true teams to shuffled team labels (Chuang & Hsu, 2023). Together, these results suggest that enhanced interbrain synchrony in cooperative compared to competitive settings could be driven by shared representations of one’s own and others’ actions, intentions, strategies, goals, etc. during cooperation, that are not present during competition.

However, these studies do not directly provide insight into what information about self and other is represented in the neural signal during competitive scenarios. Here we leverage multivariate analysis methods on electroencephalography (EEG) hyperscanning data to investigate how the human brain encodes self- and other-related information during the Rock-Paper-Scissors game to better understand the cognitive processes supporting decision performance during real-time interaction in a competitive context. Importantly, hyperscanning enabled the investigation of the processes behind winning or losing in the Rock-Paper-Scissors game, as it requires studying the cognitive and neural dynamics of the two players together in interaction. Moreover, the use of multivariate analyses on hyperscanning data allowed going beyond interbrain synchrony to more directly track neural representations related to self and other within interacting brains, consistent with the argument that understanding the neural bases of social interactions with hyperscanning requires analysis at multiple levels, between but also within brains (Zamm et al., 2024).

The Rock-Paper-Scissors game is a widely used model for studying social interaction in a competitive context, where wins, losses, or draws have equal probability for each player. The optimal strategy for both players is to be as random, and therefore unpredictable, as possible (West & Lebiere, 2001; Zhou, 2016). However, humans rarely achieve randomness (Figurska et al., 2008; Neuringer, 1986; Treisman & Faulkner, 1987) and instead, biases or strategies emerge during this game (Dyson, 2019). This includes over-selection of one option — typically Rock — adopting outcome-based strategies such as the ‘win-stay, lose-shift’ strategy, or employing cycle-based strategies, where players rotate through the options (Dyson, 2019). Humans are generally poor at being random, as evidenced by recurring patterns in the way we move, and decisions and errors we make (Figurska et al., 2008; Miyata et al., 2017; Riley & Turvey, 2002; Torre et al., 2013; Varlet & Richardson, 2015; Zhu et al., 2022), although this depends on the task and feedback (Guseva et al., 2023; Neuringer, 1986; Nickerson, 2002; Riley & Turvey, 2002; Varlet et al., 2012). By combining the Rock-Paper-Scissors game with decoding methods on EEG data, we can track the neural information about participant’s decisions, as well as the expected decisions of their opponent and information about the previous choices of self and other. Therefore, this game can serve as a useful model to investigate the neural processes underlying strategic decision-making in a competitive context that requires randomness.

Recent hyperscanning studies using fNIRS reported increased interbrain synchrony while participants played Rock-Paper-Scissors compared to rest, with effects observed in the right dorsolateral prefrontal cortex (Kayhan et al., 2022; Zhang et al., 2024), as well as the left dorsolateral prefrontal cortex and temporo-parietal junction (Kayhan et al., 2022). However, when using the more stringent comparison of playing the game alone, rather than rest, only the right temporo-parietal junction showed increased interbrain synchrony during the joint task (Kayhan et al., 2022). This suggests that observed effects in some areas could be driven largely by shared response-related processes (Burgess, 2013; Holroyd, 2022; Varlet & Grootswagers, 2024). In addition, interbrain synchrony levels did not increase when participants made explicit predictions about the outcome, aiming for either the same (cooperative condition) or a different (competitive condition) outcome as the other player, compared to when participants made implicit predictions (free play) (Kayhan et al., 2022). This suggests that interbrain synchrony was not sensitive to explicitly making predictions about the likely action of the other player and leaves open questions about the role of prediction in social decision-making contexts. As interbrain synchrony methods are activation-based, they cannot directly measure what information related to the decision or prediction is encoded in the brain. In this study, we leverage multivariate analysis methods as a powerful alternative (Moerel et al., 2025; Varlet & Grootswagers, 2024; Zada et al., 2024), combining these methods with the high temporal resolution of EEG hyperscanning to specifically capture self-other decision-related information within each player’s brain at different task stages during competitive social interactions. Specifically, for each participant, we assess the time-course of the presence of neural information about 1) the response decision of the participant, 2) the prediction about the likely response of the opponent, and 3) the knowledge about previous events, related to both self and other. Furthermore, these neural representations were compared between overall match winners and losers to understand how they might drive success.

We tested 31 pairs of participants playing a computerised version of the classic Rock-Paper-Scissors game. We recorded 64-channel EEG, while each pair played 480 games total. Behavioural results corroborated previous research, showing that participant’s responses were not fully random, even though randomness is the best strategy in this competitive game (Glover & Dixon, 2017; West & Lebiere, 2001; Zhou, 2016). Multivariate decoding methods on the EEG signals revealed the neural encoding of players’ own decisions in the current trial. Additionally, information about the participant’s own decision on the previous trial, as well as that of the opponent, was also encoded in the neural signal – but only for individuals who lost the overall match, suggesting that maintaining prior-trial information could have potentially hindered performance. These findings provide new insights into self-other neural processes supporting decision-making and the influence of cognitive biases and priors in competitive social settings.

## 2. RESULTS

### 2.1. Behavioural strategies and biases

We continuously recorded 64-channel EEG data from 62 participants, grouped into 31 pairs, using the BioSemi Active-Two electrode system (BioSemi, Amsterdam, The Netherlands). Participants were seated at a computer in separate rooms and played 480 games of a computerised version of the Rock-Paper-Scissors game. In each game, both players choose between Rock, Paper, or Scissors. The outcome of the game was determined by three rules (Figure 1A): Rock beats Scissors, Scissors beat Paper, and Paper beats Rock. If both players choose the same response, the game is a tie. For each player, wins, losses, or draws therefore have equal probability. An overview of the experiment is shown in Figure 1B. Each game consisted of three phases: Decision (2 s), Response (2 s) and Feedback (1 s), which allowed a stage-wise analysis of information in the EEG responses time-locked to each stage without interference from future stages (Moerel et al., 2024). During the Decision phase, the participants could decide which response to select. During the Response phase, participants selected their response (Rock, Paper, or Scissors), and during the Feedback phase, the outcome of the game was displayed.

**Figure 1.**
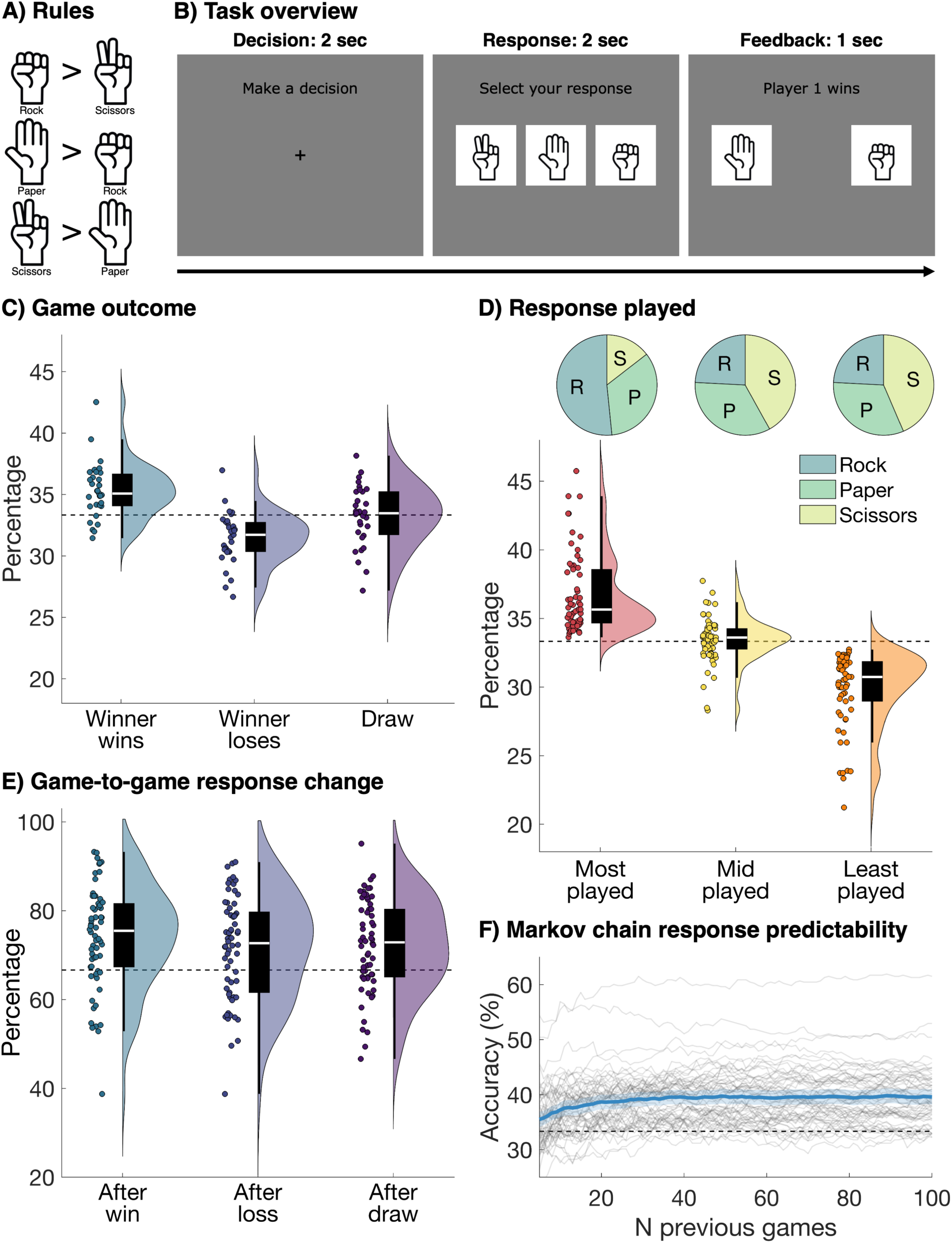
Task overview and behavioural responses. **A)** The rules of the game as shown to the participants. Rock beats Scissors, Scissors beat Paper, and Paper beats Rock. The three options are displayed as a cartoon of the hand shapes associated with the game. Rock was displayed as a fist, Paper as an open hand, and Scissors as a fist with the index and middle fingers extended. Images were sourced online from www.dreamstime.com. **B)** Each game consisted of three phases. In the Decision phase (left panel), participants saw a fixation cross with the prompt to “Make a decision” for 2 seconds. In the Response phase (middle panel), participants saw the three options with a prompt to “Select your response”. The order of the three options was randomised for each participant and block. Participants used the buttons on a button box to select their response. In the Feedback phase (right panel), participants received feedback about the response of player 1 on the left and player 2 on the right. Feedback about the outcome was displayed above as “Player 1 wins”, “Player 2 wins”, or “Draw”. The feedback was shown for 1 second. **C)** The distribution of the three possible game outcomes over pairs: the overall match winner wins (left), the overall match winner loses (middle), and draw (right). The white line on the box plot shows the median, each dot shows one pair. Theoretical chance is 33.3%. **D)** The raincloud plots show the distribution of how often the most, mid, and least chosen responses are played. The closer to 33.3%, the less bias there is. The dots show individual participants instead of pairs. The pie charts above show for each response frequency (most, mid, and least chosen) the proportion of Rock, Paper, and Scissors responses: 51.61% of participants had Rock as their most played response, followed by Paper (33.87%), and Scissors (14.52%). **E)** The percentage of games where the response changes across two consecutive games, split by the outcome of the previous game: after a win (left), loss (middle), or draw (right). All plotting conventions are the same as in Figure 1C, but theoretical chance is 66.7%. **F)** The accuracy of a Markov chain in predicting which response a player will make. This can be used as a measure of how predictable the responses of the participants are. The x-axis shows the different window sizes of the Markov chain. Grey lines show individual participants, the thick blue line shows the group average and the shaded area around the group average shows the 95% confidence interval. Theoretical chance is 33.3%.

The behavioural results revealed that selected responses were not completely random and that biases and strategies across players varied. Figure 1C shows the distribution of the three possible game outcomes for each pair: the overall match winner wins the game, the overall match winner loses the game, and the game is a tie. The overall match winner is defined as the person with the greatest number of wins out of the full 480 games. The distribution across pairs shows that although there was a clear winner for some pairs, the overall match winner and loser performed very similar in many pairs. Figure 1D shows the distribution of how often the most, mid, and least chosen responses are played. The closer to 33.3%, the less bias there was. The data illustrate that there was a spread in participants’ bias towards one option. The pie charts in Figure 1D show separately for the most, mid, and least chosen responses the proportion of *Rock*, *Paper*, and *Scissors* responses. In line with previous work (Dyson et al., 2016; Forder & Dyson, 2016; Wang et al., 2014; Xu et al., 2013), there was a bias towards Rock, 51.61% of participants had Rock as their most played response, followed by Paper (33.87%). Only 14.52% of participants had Scissors as their most played response. We also assessed the change between two consecutive responses, with a completely random player having a 33.3% chance of playing the same response twice in a row. Figure 1E shows the chance of a game-to-game response change, split by the outcome of the previous game. Many participants were biased towards changing their response, regardless of the outcome of the previous game. Finally Figure 1F shows the predictability of each player, as obtained via a Markov chain using various number of previous games. An accuracy of 33.3% means the responses of the player are completely unpredictable, which would be the best strategy. The data show that most participants were not completely unpredictable, as the Markov chain accuracy was above chance. These findings highlight a variation in participants’ strategies and responses, revealing individual biases.

### 2.2. Neural decoding of player and opponent decisions

We used linear discriminant analysis classifiers to investigate if there was information in the EEG signal about 1) the response made by the player, 2) the response made by the other player, 3) the player’s response in the prior trial, and 4) the response of the other player in the prior trial. We did this separately for each 250 ms time bin and each participant using a 10-fold cross validation (Grootswagers et al., 2017). We divided the data into 10 folds (i.e. subsets), trained the classifier using the data from all-but-one subset, and tested on the left-out subset. We repeated this 10 times, leaving out a different subset each time. We removed no-response trials and created 20 pseudo trials for each fold and response by averaging 4 trials belonging to the same fold and response type before decoding to enhance the signal to noise ratio (Grootswagers et al., 2017; Scrivener et al., 2023). For the channel searchlight, we repeated the same analysis for each individual channel and 4 to 5 neighbouring channels, to obtain topographies of the decoding accuracy. The results are shown in Figure 2 and provide evidence that information about the player’s own response was encoded in the EEG signal during the Decision phase (max BF = 57) as well as the Response phase (max BF = 729,735) and the Feedback phase (max BF = 16,028), as indicated by the Bayes Factors. There was no information about the other player’s response during the Decision and Response phases, suggesting the players were not able to determine the likely response of their opponent above chance. However, the EEG signal contained information about the other player’s response during the Feedback phase (max BF = 87,847), likely reflecting the visually presented feedback about the response of their opponent. In addition, there was evidence that the player’s own previous response was encoded during the Decision phase (max BF = 8), and anecdotal evidence during the Response phase (max BF = 4), suggesting that information about the previous response could be used during the decision-making on the current trials. Finally, there was information about the other player’s previous response during the Decision phase (max BF = 16,659), suggesting this information was incorporated in the decision about which move to play. These results show the data contain information about the player’s current and previous decisions, as well as information about the previous decisions of the other player. Moreover, the topographies show that the encoding of current and previous decisions during the Decision and Response phases is driven by distributed responses across the brain. This contrasts with the Feedback phase, where we see a strong contribution of posterior channels to the above chance decoding, in line with more visually driven responses, as participants receive visually presented feedback about their own, as well as their opponent’s response. These spatial distributions are in line with the decoding accuracies in the Decision and Response phases being driven by decision-related information.

**Figure 2.**
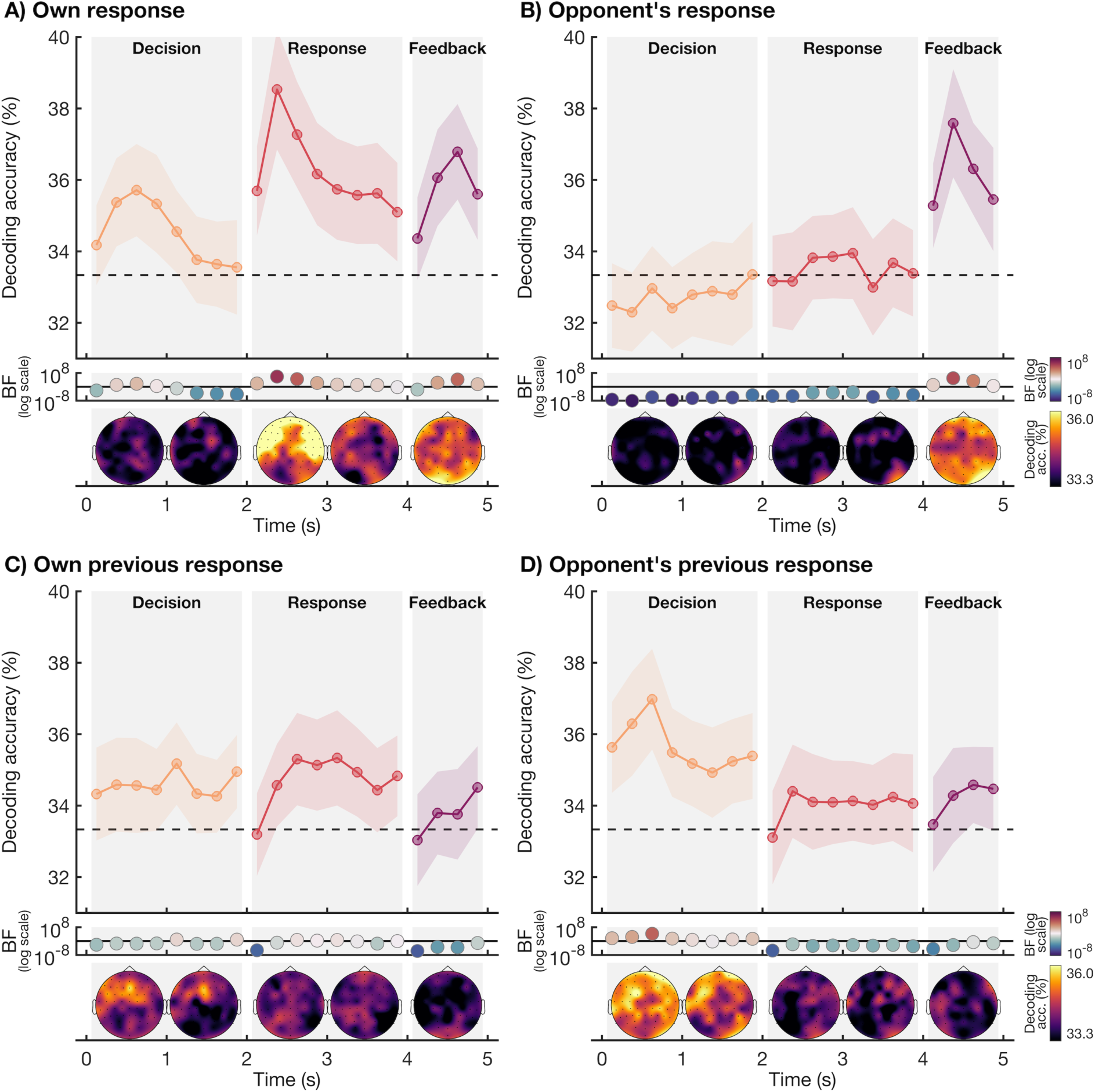
EEG decoding accuracy of the player’s own responses and those of their opponent. The plotting conventions are the same for all four plots showing the classification accuracy over time for the three phases: Decision (0 – 2 s, orange), Response (2 – 4 s, red), and Feedback (4 – 5 s, purple). Decoding accuracies are obtained for 250 ms time bins. We used a linear classifier to determine whether there was information in the pattern across the EEG channels about **A)** the response the player chose **B)** the response played by the opponent, **C)** the response the player chose on the previous trial, and **D)** the response the opponent chose on the previous trial. The Bayes Factors are shown below the plot on a logarithmic scale. Bayes Factors below 1 are plotted in blue colours and show evidence for chance decoding, and Bayes Factors above 1 are plotted in red colours and show evidence for above chance decoding, with the colour intensity reflecting the evidence for either hypothesis. The topographies, based on a channel searchlight, are shown below the Bayes Factors. We obtained a topography for each 250 ms time bin in the decoding and then collapsed across 1-second time bins. Lighter colours reflect higher decoding accuracies.

To assess whether the information that was present in the brain was modulated by whether the player won or lost over the entire experiment, we split the decoding accuracies for the overall match winners and losers (Figure 3). The player’s own response was encoded in the EEG signal during the Response and Feedback phases for both winners (max BF = 573) and losers (max BF = 3,337). Both groups showed neural information about the other player’s response during the Feedback phase only. Importantly, the results indicated that only losers showed neural encoding of their own previous response during the Response phase (max BF = 11), and of the other player’s previous response during the Decision phase (max BF = 2,382). Although inter-group Bayes Factors did not provide direct evidence of strong differences between winners and losers – indeed, most Bayes Factors showed evidence for no difference between these groups, which could be explained by the scores of the winners and losers being often very close – our results nevertheless suggest that losers encoded self and other information from previous trials, whereas winners did not.

**Figure 3.**
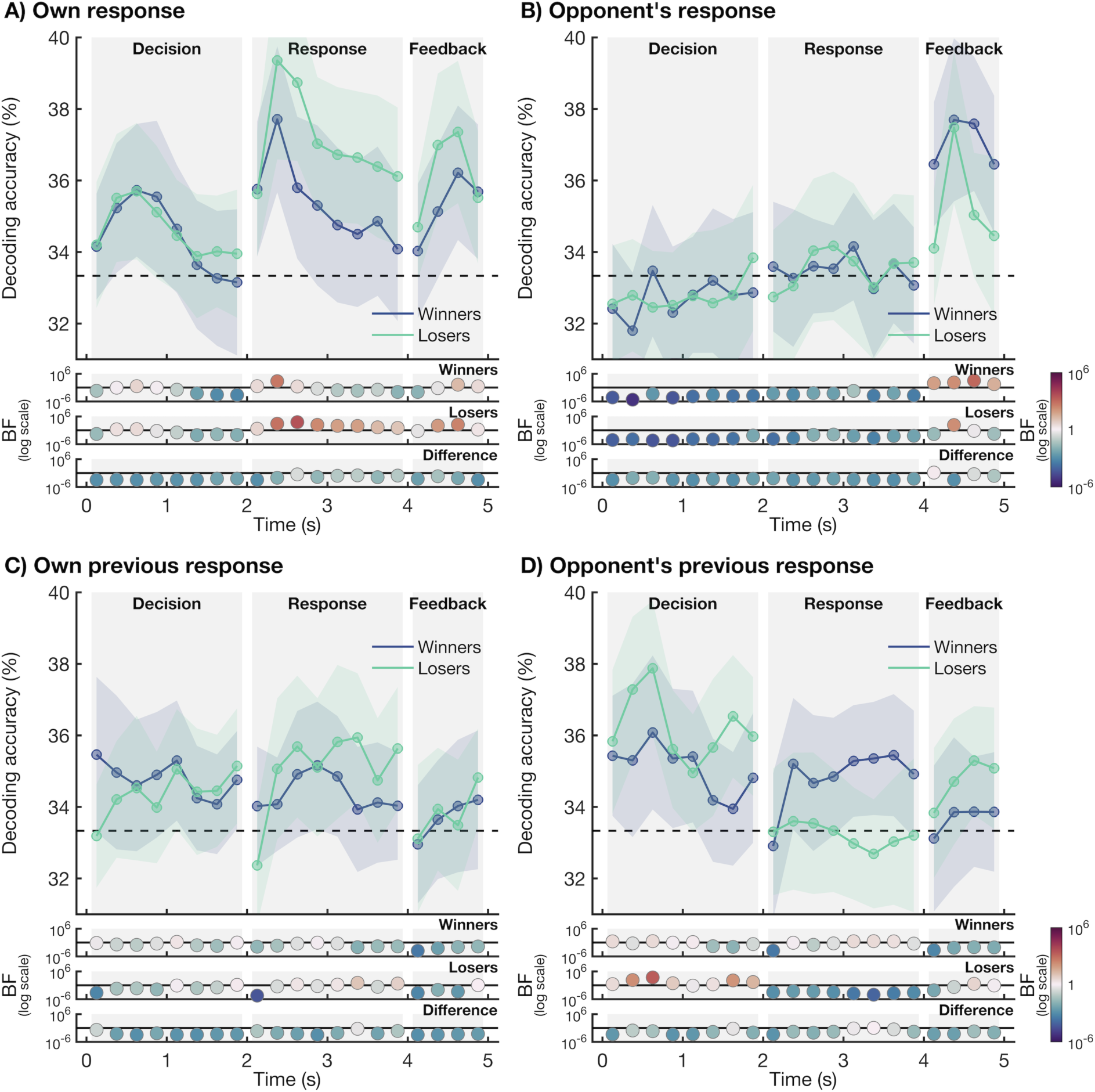
EEG decoding accuracy of the player’s own responses and those of their opponent, split by overall match winners and losers. The plotting conventions are the same as in Figure 2. We split the decoding accuracies presented in Figure 2 by overall match winners (blue) and losers (green). We determined whether there is information in the pattern across the EEG channels about **A)** the response selected by the player, **B)** the response made by the other player, **C)** the player’s response in the previous trial, and **D)** the other player’s response in the previous trial.

## 3. DISCUSSION

In this study, we investigated how the brain encodes self-other decision-related information during the competitive Rock-Paper-Scissors game. In line with previous research our participants did not act fully random, even though unpredictability is the best strategy in this competitive game (Glover & Dixon, 2017; West & Lebiere, 2001; Zhou, 2016). There were variations between players, but Rock was generally over-selected and Scissors under-selected. This finding has been widely observed in previous work (Dyson et al., 2016; Forder & Dyson, 2016; Wang et al., 2014; Xu et al., 2013). In addition, participants switched their response from trial-to-trial in a more or less predictable way, regardless of whether they won or lost on the previous trial. These behavioural biases have been observed previously in this game (Dyson, 2019), and are in line with other work that suggests people are not able to consciously generate random actions (Figurska et al., 2008; Issartel et al., 2007; Neuringer, 1986; Treisman & Faulkner, 1987).

EEG hyperscanning data revealed that there was neural information about the player’s own current response during all phases of the task, driven by the decision the participant had to make. This information was already present in the Decision phase, before the participant was asked to respond, suggesting we were able to track the decision-making of the participant as it unfolded in real-time. There was no above-chance decoding of the opponent’s decision during the Decision or Response phases. This suggests that at the level of the group, participants could not reliably predict the next move of their opponent. This finding fits with the previous hyperscanning literature, that found weaker interbrain synchrony during competition compared to cooperation (Cui et al., 2012; Lu et al., 2019; Sinha et al., 2016), as well as evidence for interbrain coupling during cooperation and interbrain decoupling during competition when comparing true teams to shuffled team labels (Chuang & Hsu, 2023). This suggests that during cooperation, participants may form a shared representation of goals, intentions, or actions of self and other, facilitated by behavioural predictability. However, during competition, behavioural unpredictability is advantageous (Glover & Dixon, 2017; Vesper et al., 2011), potentially driving a reduction in shared representations and therefore interbrain synchrony. Our results indeed showed that participants were too unpredictable for their opponents to reliably encode their next move. Finally, above chance decoding of the previous responses of self and other show that the brain actively encodes knowledge about previous trials during the Decision phase of the current trial. In determining their response on the current game, participants likely rely on what their opponent played, and to a lesser extent, what they themselves played in the previous game. This study adds to the findings from the hyperscanning literature by directly assessing the content of the neural representations about self and other, enabling us to distinguish between neural information about decisions, expectations about the opponent, and knowledge about previous interactions.

To gain insight into the neural representations that may underlie successful versus unsuccessful competitive behaviour, we tracked self- and other-related neural representations within the brains of winners compared to losers. While decoding results split by overall match winners and losers showed no differences in brain coding of decision-related information, we did find a different signature of decoding results between the two groups. Small differences in wins and losses in some pairs suggest that chance, not strategy, determined the winner, likely adding noise and obscuring potential group differences. Specifically, we found that whereas we could track the decision-making on the current trial for both the winners and losers, only the overall match losers encoded information about what they played themselves and what the other player chose on the previous trial. This suggests that losers likely incorporated information about the previous game, whereas this was not the case for the overall match winners. This reliance on previous responses, both of self and other, might hinder these participants, as the best strategy is to be as random, and therefore unpredictable, as possible (Glover & Dixon, 2017; West & Lebiere, 2001; Zhou, 2016). By comparing the individual neural representations of winners and losers, rather than focusing on interbrain synchrony, this study provides more direct insight into which self- and other-related representations are associated with a competitive advantage.

Previous work on the Rock-Paper-Scissors game has used fNIRS to investigate prediction-related processes (Kayhan et al., 2022; Zhang et al., 2024). Although the results showed increased interbrain synchrony during the interactive task compared to playing the game alone, this measure did not increase when making explicit predictions (Kayhan et al., 2022). Previous work has also found increased interbrain synchrony in various regions when playing the game compared to rest (Kayhan et al., 2022; Zhang et al., 2024), but these findings could be driven by shared response-related processes (Burgess, 2013; Holroyd, 2022; Varlet & Grootswagers, 2024). In a recent study, Varlet and Grootswagers (2024) showed that interbrain synchrony measures are not sensitive to the information contained in the neural data. To address this, they introduced Interbrain RSA, a multivariate analysis method designed to assess the alignment of information across participants’ neural signals. Building on this, we used decoding, a complementary multivariate approach (Grootswagers et al., 2017), to show that we can track the decision-related information about the responses of the participant and their opponent at different stages in the task. While the limited response options in our design precluded the use of Interbrain RSA, we show that decoding methods are highly effective in capturing decision-related neural information, highlighting the potential of this method for hyperscanning research as an extension of Interbrain RSA (Moerel et al., 2025; Varlet & Grootswagers, 2024). Unlike interbrain synchrony measures, which often rely on control conditions where participants are at rest or do not interact, the decoding method in this study leverages direct comparisons of neural activation associated with distinct response options. This makes decoding less susceptible to confounds such as shared visual or response-related information, offering a promising method to study neural processes underpinning real-time social interactions, in both cooperative and competitive settings, including in atypical populations (Charman, 2003; Coey et al., 2012; MacRitchie et al., 2017; Moreau et al., 2024; Raffard et al., 2015; Sebanz et al., 2006; Varlet et al., 2014; Zhang et al., 2024).

In conclusion, multivariate pattern analyses of EEG hyperscanning data enabled us to reveal how the human brain encodes self-other decision-related information over time during a competitive Rock-Paper-Scissors game. Despite the optimal strategy being randomness, players displayed critical behavioural biases and predictable strategies. EEG data showed neural encoding of current decisions, with overall match losers uniquely relying on past trials, potentially hindering performance. These findings highlight the challenge of overcoming cognitive biases and reliance on prior outcomes for effective decision-making during competitive social interaction.

## 4. METHODS

### 4.1. Participants

Sixty-eight participants, grouped into 34 pairs, took part in the EEG study at Western Sydney University. Three EEG datasets were excluded from the analysis: two datasets had an issue with CMS which resulted in no EEG data for one of the players, and another dataset did not contain triggers. The sample included in this dataset consisted of 31 pairs (62 participants). Participants in this sample were 38 females / 22 males / 2 non-binary, 55 right-handed / 6 left-handed / 1 ambidextrous, mean age 27.85 years, SD = 6.73 years, range 18 – 47 years. All participants reported normal or corrected-to-normal vision. The EEG session took approximately 2 hours in total to complete, and participants received a payment of $60 AUD. The study was approved by the Human Ethics Committee of the University of Western Sydney, and all participants provided written informed consent (H13092).

### 4.2. Stimuli and experiment

During the experiment, the two participants in each pair played a computerised version of the competitive Rock-Paper-Scissors game. Participants were seated behind a computer screen in separate rooms. In each game, both players choose between Rock, Paper, or Scissors. The outcome of the game is determined by three rules (Figure 1B): Rock beats Scissors, Scissors beat Paper, and Paper beats Rock. If both players choose the same response, the game is a tie. For each player, wins, losses, or draws therefore have equal probability.

An overview of the experiment is shown in Figure 1C. Each game consisted of three phases: Decision (2 s), Response (2 s) and Feedback (1 s), allowing us to assess decision-related information in the EEG signal at each stage without interference from future stages (Moerel et al., 2024). The Decision screen consisted of a central fixation cross and a prompt to “Make a decision”. During this phase, the participants could decide which response to select. During the Response phase, participants saw a prompt to “Select your response”, and the three options were displayed as a cartoon of the hand shapes associated with the game (Figure 1A); Rock was displayed as a fist, Paper as an open hand, and Scissors as a fist with the index and middle fingers extended, forming a V. The order of the Rock, Paper and Scissors images was chosen at random for each participant and each block but stayed the same for all games within a block. During this phase, participants used a 3-button box to select their response. The order of the three stimuli on the screen indicated the mapping between the buttons and responses, and the response mapping therefore changed between blocks. This means that taking together the data from the whole experiment, the EEG signal was minimally contaminated by motor preparation and/or execution signals. The Response screen automatically timed out after 2 s, even if no response was given by one or both of the participants. The Response phase was followed by the Feedback phase, where the outcome of the game was displayed for 1 s. Participants saw the response of player 1 on the left of the screen, and the response of player 2 on the right of the screen. If no response was given, “No Response” was displayed. The outcome was displayed above the response images, as “Player 1 wins”, “Player 2 wins”, or “Draw”. If one of the participants did not respond, the other participant won the game.

The experiment consisted of 480 games of Rock-Paper-Scissors, divided into 12 blocks of 40 games, and took approximately 45 minutes to complete. The experiment was displayed on a mid-grey background using a VIEWPixx monitor at 120 Hz, using custom C++ code. Participants were seated approximately 60 cm from the screen. The images were displayed at 324 by 324 pixels (approximate 8.20 by 8.20 degrees of visual angle) and the fixation cross was 54 by 54 pixels (approximate 1.37 by 1.37 degrees of visual angle).

### 4.3. EEG acquisition and pre-processing

We collected continuous EEG data from 64 channels, digitised at a sampling rate of 2048 Hz, using the BioSemi Active-Two electrode system (BioSemi, Amsterdam, The Netherlands). Electrode placement followed the international 10-20 system (Oostenveld & Praamstra, 2001). The dataset consists of the raw EEG data as well as pre-processed data. We applied the following pre-processing steps, using the FieldTrip toolbox (version 20240110) in MATLAB (Oostenveld et al., 2011). First, we re-referenced the data to the common average, and epoched the data from 200 ms before the onset of the Decision screen to 5000 ms after the onset of the Decision screen. The onset of each trial, in seconds as well as samples, can be found in the events.tsv file. We did not apply filtering, as this has been shown to cause artefacts or temporally smear the signal (Delorme, 2023; Grootswagers et al., 2017; van Driel et al., 2021). We identified noisy channels through visual inspection, which can also be found in the participants.tsv file. We interpolated noisy channels based on neighbouring channels, using the *ft_channelrepair* function with a distance measure of 0.5 cm. We then down-sampled the data to 256 Hz. Finally, we made three separate epochs for each trial, locked to the onset of the Decision screen (−200 ms – 2000 ms), Response screen (−200 ms – 2000 ms), and Feedback screen (−200 ms – 1000 ms) respectively. We applied baseline corrections for each separate epoch, using the window from −200 m to 0 ms, locked to the screen onset. We then averaged the resulting data into 250 ms time bins, resulting in a total of 20 time bins for the 0 to 5000 ms time-course.

### 4.4. Behavioural analysis

To examine the behavioural responses recorded during the EEG session, we performed the following analyses. To assess the difference in performance between winners and losers, we obtained the distribution of game outcomes across pairs, categorising results as wins for the overall match winner, wins for the overall match loser, or ties. We examined the response bias by analysing how frequently the most, mid, and least chosen responses were played. To explore this further, we looked at the proportion of Rock, Paper, and Scissors responses separately for the most, mid, and least chosen responses. Additionally, we assessed the distribution of response changes between consecutive responses, split by the outcome of the previous game. Finally, we used a Markov chain model (Norris, 1998) with varying window sizes (5 to 100 previous games, in steps of 1) to predict each player’s most likely response. We then calculated the prediction accuracy as a measure of response predictability for each participant. Together, these analyses provide insight into the strategies, response biases, and predictability of the participants.

### 4.5. Decoding analysis

We used a time-varying brain decoding analysis (Grootswagers et al., 2017) implemented using the CoSMoMVPA toolbox (version 1.1.0) (Oosterhof et al., 2016) to determine whether there was information about the response decisions made by the player and opponent, based on the raw activation values across the 64 EEG channels. Specifically, we used a regularised (λ = 0.01) linear discriminant analysis (LDA) classifier to determine if there was information about 1) the response made by the player, 2) the response made by the other player, 3) the player’s response in the prior trial, and 4) the response of the other player in the prior trial. We opted for this classifier as it provides a good balance between performance and computational requirements (Grootswagers et al., 2017; Guggenmos et al., 2018). Before decoding, we removed no-response trials and made 20 pseudo trials for each fold and response (Rock, Paper, or Scissors) by averaging 4 trials of the same fold and response to enhance the signal to noise ratio (Grootswagers et al., 2017; Scrivener et al., 2023). We then trained a linear classifier to distinguish between the different classes of interest (e.g. the different responses made by the player) based on the pattern of activation across all 64 EEG channels. We used the EEG channel voltages, separate for each trial and time bin, as features for the decoding. We did this separately for each individual participant. We used a 10-fold cross validation, where we split the dataset into 10 parts (i.e. folds). We trained the linear classifier on the data from 9 of those folds and then tested the classifier performance on the left-out fold. We did this in an iterative fashion, where each fold was left out as a test fold once. We then calculated the classifier accuracy, separately for each individual participant and time bin, by determining the percentage of times the classifier predicted the correct class (e.g. response is rock). We calculated the decoding accuracy separately for each of the 3 possible responses and then averaged these measures to obtain the decoding accuracy for each time bin. If the classification accuracy is higher than would be predicted by chance, we can say that there is information in the neural response about the condition that is decoded. To ensure that motor signals were not a reliable signal for the classifier, distinguishing between the different response options during the Response phase, we changed the response button mapping between blocks. A given response option (e.g. Rock) was therefore associated with all response buttons in the training data. In addition, the mapping between the responses and buttons differed between the participants in the dyad, which means any similarity in responses between individuals in the dyad would not translate into similarities in motor preparation or execution. Together, these measures eliminated the chance of motor contamination in the decoding. For the channel searchlight, we repeated the same analysis above for individual clusters of channels. Each cluster consisted of the main channel and 4 or 5 neighbouring channels.

### 4.6. Statistics

To determine whether there was evidence for above chance decoding of decision-related information, we used Bayesian statistics (Dienes, 2011; Kass & Raftery, 1995; Morey et al., 2016; Rouder et al., 2009; Wagenmakers, 2007), with the Bayes Factor R package (Morey & Rouder, 2018). To calculate the Bayes Factors, we applied Bayesian directional t-tests for each time bin. We used the method described by Teichmann and colleagues (2022), using a null interval between d = 0 and d = 0.5 to exclude small effect sizes. To calculate Bayes Factors for the difference between the overall match winners and losers, we used a two-tailed t-test.

## 5. CODE AND DATA AVAILABILITY

All analysis codes, figures, and an example dataset are publicly available via the Open Science Framework (https://doi.org/10.17605/OSF.IO/YJXKN). The data are named and organised according to the BIDS standard (Gorgolewski et al., 2016; Pernet et al., 2019). The full dataset will be made publicly available upon publication.

## 6. ACKNOWLEDGMENTS

This research was supported by an Australian Research Council Discovery Project DP220103047 (M.V.), an Australian Research Council Discovery Early Career Researcher Award DE230100380 (T.G.), and H2020 Marie Skłodowska-Curie grant (F.C.; agreement no. 893960).

## Notes

https://doi.org/10.17605/OSF.IO/YJXKN

### Competing Interest Statement

The authors have declared no competing interest.

### Summary of Updates

We have provided additional details about the decoding analysis.

